# VeChat: Correcting errors in long reads using variation graphs

**DOI:** 10.1101/2022.01.30.478352

**Authors:** Xiao Luo, Xiongbin Kang, Alexander Schönhuth

## Abstract

Error correction is the canonical first step in long-read sequencing data analysis. The current standard is to make use of a consensus sequence as a template. However, in mixed samples, such as metagenomes or organisms of higher ploidy, consensus induced biases can mask true variants affecting haplotypes of lower frequencies, because they are mistaken as errors.

The novelty presented here is to use graph based, instead of sequence based consensus as a template for identifying errors. The advantage is that graph based reference systems also capture variants of lower frequencies, so do not mistakenly mask them as errors. We present VeChat, as a novel approach to implement this idea: VeChat distinguishes errors from haplotype-specific true variants based on variation graphs, which reflect a popular type of data structure for pangenome reference systems. Upon initial construction of an ad-hoc variation graph from the raw input reads, nodes and edges that are due to errors are pruned from that graph by way of an iterative procedure that is based on principles from frequent itemset mining. Upon termination, the graph exclusively contains nodes and edges reflecting true sequential phenomena. Final re-alignments of the raw reads indicate where and how reads need to be corrected.

Extensive benchmarking experiments demonstrate that PacBio and ONT reads corrected by VeChat contain 4 to 15, or, respectively, 2 to 10 times less errors than when corrected state of the art approaches. VeChat is implemented in an easy-to-use open-source tool and publicly available at https://github.com/HaploKit/vechat.

## Background

Third generation sequencing (TGS) such as single-molecule real-time (Pacific Biosciences, or short PacBio) or nanopore sequencing (Oxford Nanopore Technologies or short ONT) has been emerging rapidly over the last few years. Beyond the obvious reason that TGS reads tend to be longer by orders of magnitude—read length ranges from several kbp up to even a few Mbp (Logsdon *et al.*, 2020), whereas next-generation sequencing reads usually span only a few hundred base pairs—the fact that TGS is relatively inexpensive adds to its popularity. Moreover, TGS circumvents polymerase chain reaction (PCR) as part of its protocol, which prevents related biases. Thanks to these advantages, TGS has been able to make decisive contributions in various areas of application. Prominent examples are haplotype phasing (Schrinner *et al.*, 2020), genome assembly (Jain *et al.*, 2018b; Ruan and Li, 2020; Shafin *et al.*, 2020; Kolmogorov *et al.*, 2020; Miga *et al.*, 2020) and (complex) variant calling (Edge and Bansal, 2019; Thibodeau *et al.*, 2020; Fujimoto *et al.*, 2021).

On the other hand, however, the downside of TGS are the significantly elevated error rates the reads are affected with. For example, PacBio CLR and ONT reads, as the currently most representative examples of TGS reads, contain 5% to 15% errors (Logsdon *et al.*, 2020). This comes in obvious contrast to NGS short reads, whose error rates usually do not exceed 1%. The fact that the majority of errors affecting long reads consists of insertions and deletions adds to the difficulties because it prevents the application of principles and straightforward adaptation of tools for correcting errors in short reads. At any rate, direct usage of raw TGS reads is hardly possible in the majority of relevant applications. So, novel methods and tools are required.

Because correcting errors in TGS reads is imperative for sound analyses and because correcting errors in TGS reads is a must, various TGS read error correction methods have been presented in the meantime. The corresponding range of methods can be divided into two major categories: hybrid correction and self correction. While hybrid correction addresses to integrate short reads in the process, self correction seeks to correct errors without auxiliary data.

Clearly, hybrid correction reflects a sound and reasonable approach in general (see (Hackl *et al.*, 2014; Salmela and Rivals, 2014; Firtina *et al.*, 2018; Morisse *et al.*, 2018) for prominent approaches). However, hybrid correction suffers from some pragmatic issues. First, while long reads can span repetitive regions, short reads cannot; this introduces ambiguities in the process of assigning short to long reads (or vice versa), and as a consequence biases in the quality of the correction. Second, short reads re-introduce PCR induced biases. For example, certain areas of genomes are not sufficiently covered by short reads because of sequence content (e.g. GC content). This hampers error correction in these areas. Last but not least, sequencing genome samples employing several different protocols can be entirely impossible, which prevents the application of hybrid error correction in the first place.

Self-correction, as the second class of methods, does not suffer from any of these issues. However, because of the lack of external (e.g. short read based) assistance, self-correction faces other methodically principled challenges. It is key to overcome these challenges before one can profit from the great practical advantages of self error correction.

In terms of prior, related work, self-correction can be further divided into three sub-categories, each of which is characterized by particular algorithmic strategies and methodical foundations.

The first, and most common of the three categories is based on multiple sequence alignments (MSAs). For prior approaches and tools that crucially rely on computing MSAs, see Racon (Vaser *et al.*, 2017), the error correction module of the assembler Canu (Koren *et al.*, 2017), and FLAS (Bao *et al.*, 2019). The second principled class of approaches relies on de Bruijn graphs (DBGs). Corresponding tools employ DBGs at some point crucial for the correction process. The prevalent tool to consider is Daccord (Tischler and Myers, 2017), which is based on raising local DBGs, where local refers to DBGs reflecting relatively small segments of the genome the reads stem from.

The third class of self correction methods collects approaches that make combined use of both MSAs and DBGs during the process. Such methodically combined approaches seek to balance the advantages and disadvantages of the two concepts, MSAs on the one hand, and DBGs on the other hand. Prominent tools that make successful, combined use of MSAs with DBGs are LoRMA (Salmela *et al.*, 2017) and CONSENT (Morisse *et al.*, 2021).

The common denominator that unifies all of these approaches is to raise consensus sequence that reflects a summary of the reads observed, and serves as a guideline during error correction. However, sequence-based templates cannot capture ambiguities, whcih explains the corresponding biases during the correction process: for each polymorphic site, one needs to decide on one of the possible alleles while discarding all others. As a consequence, these approaches tend to mask variation that characterizes little covered haplotypes or strains in mixed samples (metagenomes, cancer genomes) or polyploid genomes. Because the haplotypes affected virtually disappear, downstream analyses remain blind with respect to them.

To address this issue, we developed VeChat, a self-correction method to perform *haplotype-aware error correction* for long reads. From a larger perspective, VeChat considers the full spectrum of all possible haplotypes that possibly affect the sample already during error correction, and not only thereafter. This reflects a novelty for ploidies larger than two, because earlier approaches only deal with diploid scenarios (Luo *et al.*, 2021). From a methodical point of view, the novelty of VeChat is to integrate variation graphs as a fundamental data structure into the process of error correction. To the best of our knowledge, VeChat is the first approach to do that.

We have tested VeChat and extensively compared it with the current state of the art on datasets reflecting various settings of current interest. Benchmarking experiments on both simulated and real data demonstrate that our approach achieves the best performance rates, across all categories of common sequencing errors.

## Results

We have designed and implemented VeChat ([V]ariation graph based [e]rror [C]orrection in [ha]plo[t]ypes), a new approach to haplotype aware long read self error correction. The key concept underlying VeChat are variation graphs. Unlike single consensus sequences, which current approaches are generally centering on, variation graphs are able to represent the genetic diversity across multiple, evolutionarily or environmentally coherent genomes. This enables to preserve haplotype specific variation during error correction also for samples of higher, known or unknown ploidy.

In this section, we first provide a high-level description of the workflow of VeChat. We then evaluate the performance of VeChat on both simulated and real data in comparison with the state of the art approaches.

### Workflow

See Figure 1 for an illustration of the workflow of VeChat. VeChat consists of two cycles. While the first cycle yields pre-corrected reads, the second cycle generates the final, corrected reads from the pre-corrected reads. Each cycle proceeds in 6 steps. While the two cycles generally agree on these 6 steps, they disagree in terms of crucial details affecting steps 1 and 4.

**Figure 1.**
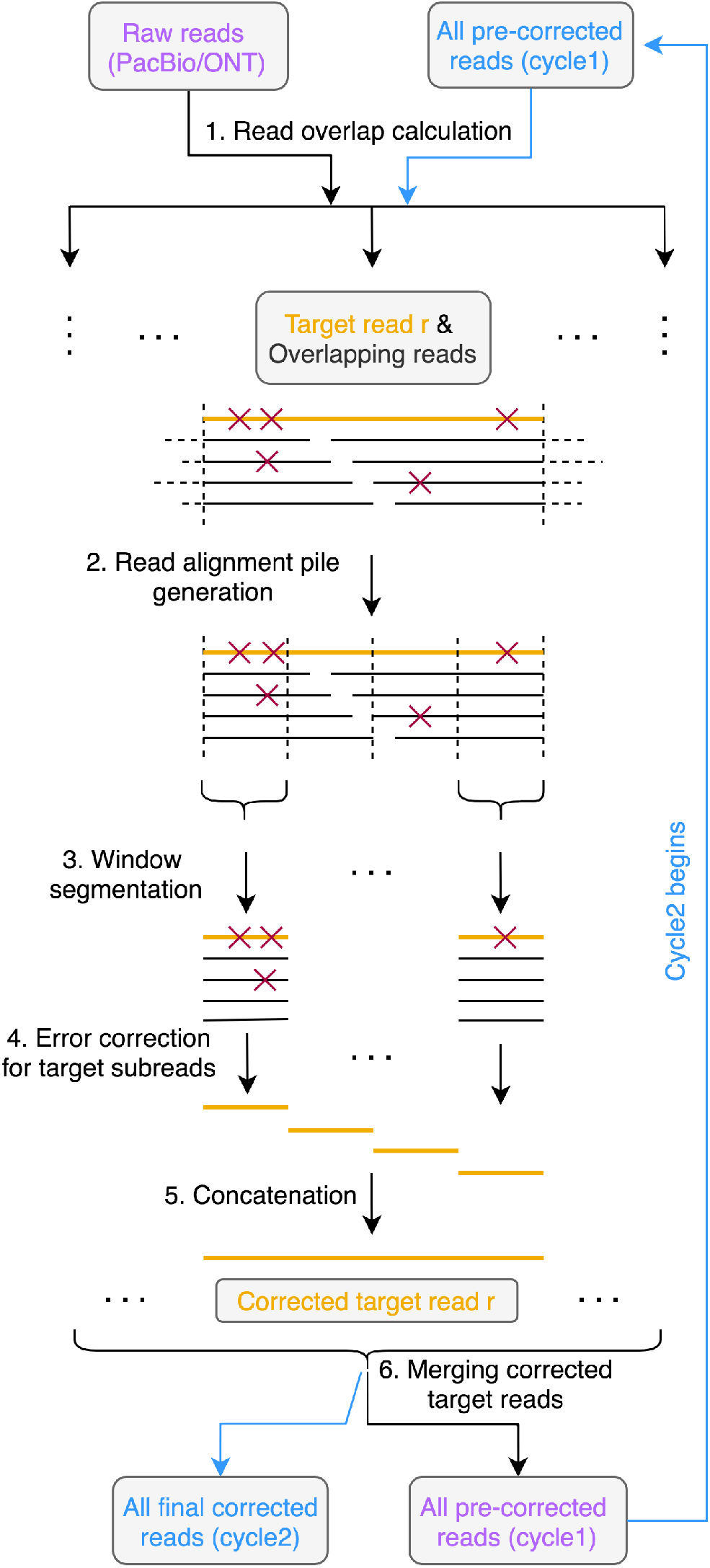
Workflow of VeChat. The input and output of cycle 1 and cycle 2 are labeled with purple and blue, respectively. Both cycle 1 and cycle 2 share the steps 1-6 except some differences in step 1 and 4. The target read is highlighted with orange. Red forks indicate the sequencing errors in reads.

During the *first cycle*, step 1 computes minimizer based all-vs-all overlaps, for which we employ Minimap2 (Li, 2018). Because using Minimap2 prevents the need for computing base-level alignments, this stage proceeds rapidly and without any further efforts.

Steps 2-6 reflect the technical core of the error correction procedure in Figure 1. In step 2, a target read is selected as the read whose errors are to be corrected, and a read alignment pile that consists of all reads that overlap it is computed. Subsequently, in step 3, the read alignment pile is divided into small segments, where each of the segments gives rise to a window like part of the pile in step 3; the part of the target read in a particular window is further referred to as ‘target subread’.

Subsequently, in step 4, the error correction for target subreads is performed in each window. Step 4 is methodically more involved, because it captures the novel, variation graph based approach; see Figure 2(a) for detailed illustrations on the version of that particular step used in the first cycle. Step 4 involves the construction of a variation graph, and pruning this graph in an iterative manner (’Graph pruning’ and ‘Graph re-pruning’ in Figure 2(a)). Pruning reflects removing spurious nodes and edges, and models nodes and edges in terms of a frequent itemset model that involves read coverage, sequencing errors and co-occurrence of characters in reads; see subsection for details. The first cycle concludes with concatenating the different ‘target subreads’ of one target read, which results in a pre-corrected read at full, original length. These pre-corrected reads then serve as input for the second cycle.

**Figure 2.**
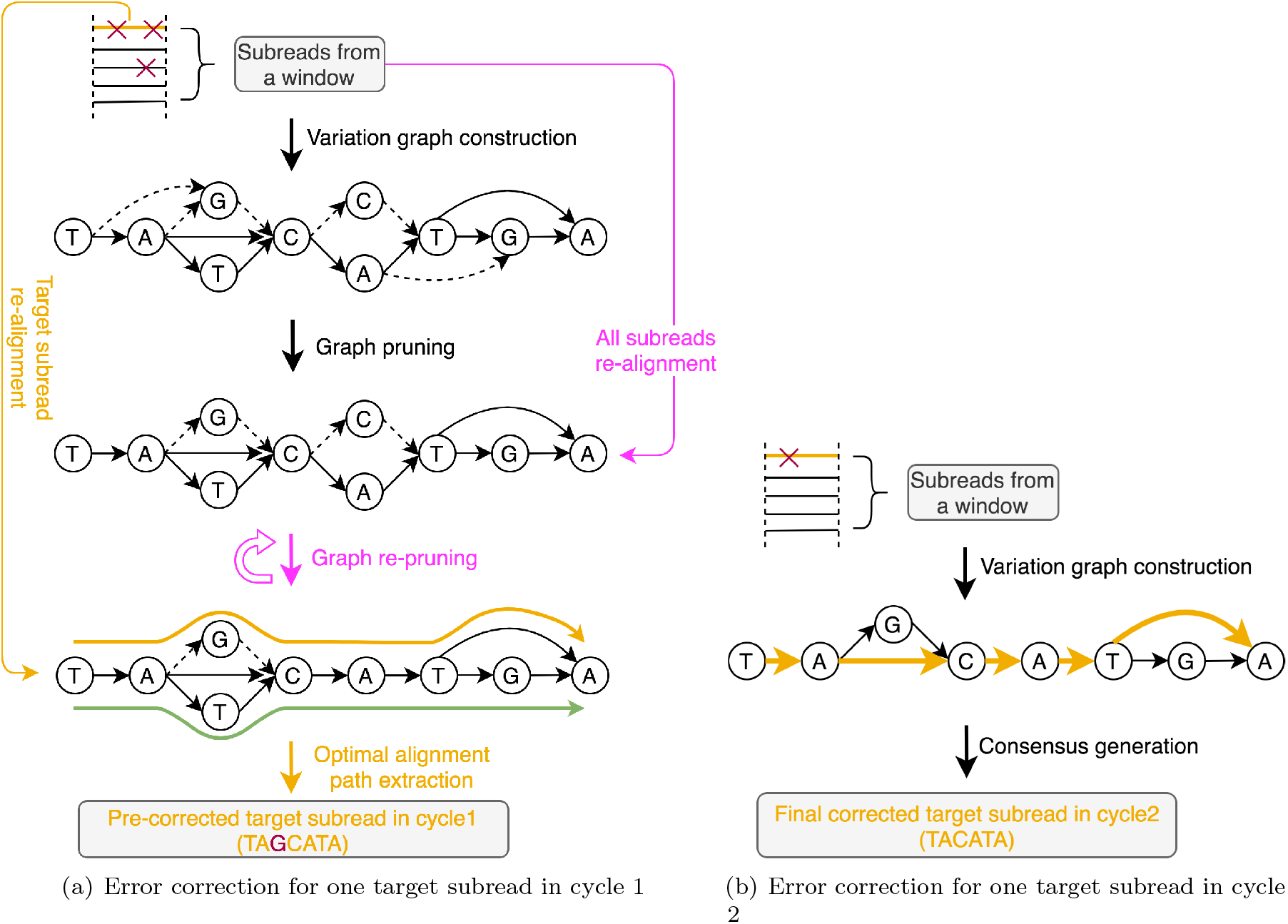
Error correction for one target subread. (a) is the error correction diagram for one target subread in cycle 1. The error correction process for the target read *r* is illustrated assuming a diploid scenario (the orange path represents the optimal alignment path, whereas the green path represents the other true haplotype). “Graph pruning” and “Graph re-pruning” refer to the core error correction procedures. These procedures rely on a variation graph that is constructed from segments of a read alignment pile that results from a multiple alignment of the target read and the reads that overlap it, see Figure 1. During graph pruning and re-pruning spurious edges (dashed arrows), induced by sequencing errors, are removed from the variation graph. The pink elements indicate that these procedures are repeated. (b) is the error correction diagram for one target subread in cycle 2. The bold orange path indicates the consensus sequence. The false nucleotide ‘G’ in the pre-corrected target subread in cycle 1 (see in (a)) is marked with red and further corrected in cycle 2.

While the (vast) majority of errors have already been corrected during the first cycle, a few errors, to be considered statistical outliers that escape correction during iterative graph pruning during the first cycle, have resisted their correction. The second cycle is supposed to spot such outliers. The second cycle is less complex than the first cycle, because it does no longer include the statistically involved graph pruning procedure; the blue elements in Figure 1 point out the different routes along which the second cycle proceeds. Overall, as above-mentioned, the second cycle is identical with the first cycle in steps 2, 3, 5 and 6. In step 1, however, beyond computing all-vs-all overlaps, base-level alignment are computed, which enables haplotype aware read overlap filtration. Step 4, as shown in Figure 2(b), then proceeds differently insofar as graph (re-)pruning is no longer part of the cycle. Instead of iterative re-pruning, which did not lead to removal of the errors that we would like to remove during this second cycle, only consensus sequences (displayed as thick yellow arrows in Figure 2(b)) are derived from the constructed variation graphs by using a dynamic programming algorithm (Lee, 2003). After concatenating the consensus sequences in step 5, joining all target reads in step 6 generates the final output of Vechat. Steps 1-6 are described in detail referring to both first and second cycle in Methods.

### Datasets

#### Simulated sequencing data

We made use of a very recent tool PBSIM2 (Ono *et al.*, 2021) to simulate PacBio CLR and Oxford Nanopore reads using built-in `P6C4` and `R103` model-based simulation profiles, respectively. Since the main application scenario of VeChat is to correct long-read sequencing data from multiple genomes, such as polyploid genomes and metagenomes, we simulated various datasets for both cases.

##### Polyploid genome data

We constructed pseudo diploid (ploidy=2, ANI: 98%), triploid (ploidy=3, ANI: 96% ~ 98%) and tetraploid (ploidy=4, ANI: 96% ~ 99%) genomes by mixing strains of *Escherichia coli* (*E. coli*) bacteria; note that Average Nucleotide Identity (ANI) is defined to measure the genome sequence similarity, which can be reported by FastANI (Jain *et al.*, 2018a), for example. All genome sequences of *E. coli* were downloaded from the NCBI database (see Supplementary Table S1 for the details of reference genomes). Reads were simulated from the haplotypes (i.e. strains) independently and upon generation mixed together to form the corresponding polyploid genome data sets (ploidy=2,3,4). We simulated both PacBio CLR and Nanopore reads for these datasets, at average sequencing coverage of 30x per haplotype and average sequencing error rate of 10%.

##### Metagenome data

Additionally, we used CAMISIM (Fritz *et al.*, 2019) to simulate two metagenomic datasets (PacBio CLR reads) of different levels of complexity. Here, we used PBSIM2 to simulate PacBio CLR reads instead of the built-in simulator in CAMISIM. The low complexity dataset consists of 10 species (20 strains), whereas the high complexity dataset consists of 30 species (100 strains). The genomes used in both datasets are derived from (Quince *et al.*, 2017) (see Supplementary Table S1 for the details). For both datasets, the average sequencing coverage of strains is about 30x and the average sequencing error rate is 10%. The relative abundances of strains range from 1.9% to 10.6% and from 0.28% to 3.3%, respectively.

#### Real sequencing data (Mock communities)

##### Yeast pseudo-diploid genome data

We constructed a pseudo-diploid genome by mixing two yeast strains (N44, CBS432) of ANI 98.4%, which are derived from Yeast Population Reference Panel (see Supplementary Table S1). The corresponding real PacBio CLR reads were downloaded from European Nucleotide Archive (ENA) under project PRJEB7245, and we only subsampled long reads with sequencing coverage of 30x per strain for further analyses.

##### NWC Metagenome data

We downloaded two real metagenomic datasets (PacBio CLR reads) derived from natural whey starter cultures (NWCs) (Somerville *et al.*, 2019) and mixed both together (see Supplementary Table S1), and then subsampled 20% reads such that we obtained a low-complexity metagenomic dataset, which contains 3 species (6 strains).

##### Microbial 10-plex Metagenome data

We downloaded raw long-read sequencing data and the corresponding reference genomes from a 10-plex multiplexed dataset which was sequenced by PacBio Sequel System, Chemistry v3.0 (https://downloads.pacbcloud.com/public/dataset/microbial_multiplex_dataset_release_SMRT_Link_v6.0.0_with_Express_2.0/). Then, we randomly subsampled 10% reads such that we obtained a mock metagenomic dataset with an average sequencing coverage about 40x, which contains 7 species (9 strains in all, ANI< 98.5%, see Supplementary Table S1).

### Benchmarking: Alternative Approaches

To enable a fair and meaningful comparison, we considered all popular state-of-the-art tools that perform TGS read self-correction. Namely, this selection includes Racon (v1.4.13) (Vaser *et al.*, 2017), CONSENT (v2.2.2) (Morisse *et al.*, 2021), Canu (v2.1.1) (Koren *et al.*, 2017) and Daccord (v0.0.18) (Tischler and Myers, 2017). We ran all tools using their default parameters.

### Metrics for evaluation

Genome assembly performance is usually evaluated by means of several commonly used metrics, as reported by QUAST V5.1.0 (Mikheenko *et al.*, 2018). See below for specific explanations, and see http://quast.sourceforge.net/docs/manual.html for full details. Note that the principled qualities of error-corrected long reads are covered by standard QUAST criteria as well; for example, low haplotype coverage reflects that true variants were mistakenly identified as errors, while error rates and misassemblies reflect that the correction procedure overlooked errors, or even confounded reads in terms of their origin through mistaken correction. As usual, corrected reads of length less than 500bp were filtered from the output before evaluation. Note that we ran QUAST with the option --ambiguity-usage one, which appropriately takes into account that our data sets reflect mixed samples (such as polyploid genomes or metagenomes).

#### Error rate (ER)

The error rate is equal to the sum of mismatch rate and indel rate when mapping the obtained corrected reads to the reference haplotype sequences.

#### Haplotype coverage (HC)

Haplotype coverage is the percentage of aligned bases in the ground truth haplotypes covered by corrected reads, which is used to measure the completeness of the corrected reads.

#### N50 and NGA50

N50 is defined as the length for which the collection of all corrected reads of that length or longer covers at least half the given sequences. NGA50 is similar to N50 but can only be calculated when the reference genome is provided. NGA50 only considers the aligned blocks (after breaking reads at misassembly events and trimming all unaligned nucleotides), which is defined as the length for which the overall size of all aligned blocks of this length or longer equals at least half of the reference haplotypes. Both N50 and NGA50 are used to assess the length distribution of corrected reads. Note that this may be of relatively little interest for corrected reads. We nevertheless display corresponding results because error correction does have an influence on these statistics.

#### Number of misassemblies (#Misassemblies)

The misassembly event in corrected reads indicates that left and right flanking sequences align to the true haplotypes with a gap or overlap of more than 1kbp, or align to different strands, or even align to different haplotypes or strains. Here, we report the total number of misassemblies in the given sequence data.

### Benchmarking results

#### Simulated polyploid genome datasets: PacBio

Table 1 shows the error correction benchmarking results for simulated PacBio CLR reads from genomes of varying ploidies, namely 2,3 and 4. VeChat achieves approximately 14 ~ 30, 9 ~ 26 and 4 11 times lower error rates on diploid, triploid and tetraploid genomes, respectively. At the same time, it maintains better or comparable performance in terms of other aspects such as number of corrected reads, completeness (haplotype coverage), number of misassemblies and length of corrected reads (as shown by N50/NGA50). In particular, VeChat drastically outperforms other tools in terms of mismatch rate (4 ~ 69 times lower than others).

**Table 1.** Error correction benchmarking results for simulated PacBio CLR reads of various polyploid genomes (ploidy=2,3,4). The average sequencing coverage per haplotype is 30x and sequencing error rate is 10%. ‘#Seq’ indicates the number of corrected reads. The error rate is equal to the sum of mismatch and indel rate. The results are sorted by the error rate in ascending order.

| Method | #Seq | Error<br>rate<br>(%) | Mismatch<br>(%) | Indel<br>(%) | Haplotype<br>coverage<br>(%) | N50<br>(bp) | NGA50<br>(bp) | # Mis-<br>assem-<br>blies |
| --- | --- | --- | --- | --- | --- | --- | --- | --- |
| **Ploidy=2** |  |  |  |  |  |  |  |  |
| VeChat | 31958 | 0.014 | 0.006 | 0.008 | 100.0 | 12556 | 38515 | 0 |
| CONSENT | 33115 | 0.194 | 0.152 | 0.042 | 99.9 | 12509 | 38318 | 38 |
| Racon | 32183 | 0.276 | 0.190 | 0.085 | 99.2 | 12514 | 38421 | 72 |
| Canu | 25924 | 0.308 | 0.183 | 0.124 | 99.9 | 13280 | 38517 | 3 |
| Daccord | 31403 | 0.423 | 0.412 | 0.011 | 99.2 | 12604 | 38598 | 3 |
| **Ploidy=3** |  |  |  |  |  |  |  |  |
| VeChat | 48085 | 0.031 | 0.015 | 0.016 | 100.0 | 12595 | 38467 | 13 |
| CONSENT | 50462 | 0.276 | 0.205 | 0.071 | 100.0 | 12511 | 38187 | 105 |
| Racon | 48986 | 0.558 | 0.427 | 0.131 | 98.7 | 12484 | 38257 | 288 |
| Canu | 37210 | 0.612 | 0.405 | 0.207 | 99.9 | 13675 | 38360 | 30 |
| Daccord | 48189 | 0.807 | 0.752 | 0.055 | 99.7 | 12485 | 38357 | 16 |
| **Ploidy=4** |  |  |  |  |  |  |  |  |
| VeChat | 62743 | 0.074 | 0.047 | 0.027 | 99.9 | 12593 | 38442 | 44 |
| CONSENT | 66315 | 0.275 | 0.180 | 0.095 | 100.0 | 12492 | 38268 | 111 |
| Racon | 64342 | 0.547 | 0.398 | 0.149 | 93.2 | 12464 | 38016 | 387 |
| Canu | 46698 | 0.549 | 0.335 | 0.214 | 99.5 | 13956 | 38255 | 86 |
| Daccord | 63440 | 0.833 | 0.790 | 0.043 | 97.5 | 12463 | 38335 | 22 |

#### Simulated polyploid genome datasets: ONT

Table 2 shows the error correction benchmarking results for simulated Oxford Nanopore reads from genomes of varying ploidy, namely 2, 3 and 4. VeChat achieves approximately 10 ~ 20, 3 ~ 9 and 2 ~ 5 times lower error rates on diploid, triploid and tetraploid genomes, respectively, while maintaining better or comparable performance in terms of all other aspects. Just as for PacBio reads, VeChat also shows better performance in terms of mismatch rate: 2 ~ 59 times lower than other correction tools, compared with indel rate.

**Table 2.** Error correction benchmarking results for simulated Oxford Nanopore reads of various polyploid genomes (ploidy=2,3,4). The average sequencing coverage per haplotype is 30x and sequencing error rate is 10%.

| Method | #Seq | Error rate (%) | Mismatch (%) | Indel (%) | Haplotype coverage (%) | N50 (bp) | NGA50 (bp) | # Mis-assemblies |
| --- | --- | --- | --- | --- | --- | --- | --- | --- |
| **Ploidy=2** |  |  |  |  |  |  |  |  |
| VeChat | 30920 | 0.022 | 0.007 | 0.014 | 99.9 | 13095 | 40612 | 5 |
| CONSENT | 32661 | 0.212 | 0.160 | 0.052 | 99.9 | 13040 | 40903 | 37 |
| Racon | 31840 | 0.346 | 0.234 | 0.112 | 99.3 | 13039 | 40725 | 120 |
| Canu | 25506 | 0.390 | 0.206 | 0.183 | 100.0 | 13820 | 40846 | 6 |
| Daccord | 31438 | 0.438 | 0.410 | 0.027 | 99.2 | 12987 | 40730 | 3 |
| **Ploidy=3** |  |  |  |  |  |  |  |  |
| VeChat | 45113 | 0.090 | 0.041 | 0.050 | 100.0 | 13130 | 40497 | 90 |
| CONSENT | 49826 | 0.298 | 0.221 | 0.077 | 99.9 | 13037 | 40877 | 103 |
| Racon | 48520 | 0.673 | 0.501 | 0.172 | 98.6 | 13028 | 39286 | 631 |
| Canu | 36962 | 0.748 | 0.453 | 0.295 | 99.9 | 14141 | 40605 | 59 |
| Daccord | 49033 | 0.821 | 0.751 | 0.069 | 99.7 | 12712 | 39193 | 11 |
| **Ploidy=4** |  |  |  |  |  |  |  |  |
| VeChat | 58739 | 0.169 | 0.098 | 0.071 | 99.7 | 13129 | 40450 | 177 |
| CONSENT | 65384 | 0.292 | 0.195 | 0.097 | 100.0 | 13033 | 40864 | 121 |
| Racon | 63670 | 0.666 | 0.469 | 0.197 | 94.9 | 13012 | 40007 | 668 |
| Daccord | 65072 | 0.840 | 0.784 | 0.056 | 96.9 | 12606 | 39076 | 22 |

#### Simulated metagenome datasets

Table 3 shows the error correction benchmarking results for simulated PacBio CLR reads of metagenomic datasets with different complexity. VeChat achieves approximately 6 ~ 7 and 3 ~ 4 times lower error rates on low and high complexity metagenomes, respectively, while maintaining comparable performance in terms of other aspects. Particularly, VeChat drastically outperforms other tools in terms of mismatch rate (6 ~ 12 times lower) on the low complexity dataset.

**Table 3.** Error correction benchmarking results for simulated PacBio CLR reads of metagenomic datasets with different complexity. The average sequencing coverage of strains is about 30x and the sequencing error rate is 10%.

| Method | #Seq | Error rate (%) | Mismatch (%) | Indel (%) | Haplotype coverage (%) | N50 (bp) | NGA50 (bp) | # Mis-assemblies |
| --- | --- | --- | --- | --- | --- | --- | --- | --- |
| **Low complexity (20 genomes)** |  |  |  |  |  |  |  |  |
| VeChat | 293466 | 0.036 | 0.020 | 0.015 | 96.9 | 11866 | 29555 | 104 |
| Racon | 299053 | 0.200 | 0.122 | 0.078 | 91.7 | 11811 | 29514 | 794 |
| CONSENT | 299333 | 0.214 | 0.149 | 0.065 | 98.4 | 11841 | 29556 | 515 |
| Canu | 253381 | 0.259 | 0.134 | 0.125 | 97.4 | 12370 | 29457 | 139 |
| Daccord | 298284 | 0.259 | 0.243 | 0.016 | 92.8 | 11862 | 29595 | 280 |
| **High complexity (100 genomes)** |  |  |  |  |  |  |  |  |
| VeChat | 1441190 | 0.088 | 0.061 | 0.026 | 97.5 | 11886 | 30129 | 2774 |
| CONSENT | 1497216 | 0.274 | 0.163 | 0.112 | 99.4 | 11839 | 30204 | 3263 |
| Canu | 1185152 | 0.354 | 0.192 | 0.162 | 99.0 | 12706 | 30016 | 873 |
| Racon | - | - | - | - | - | - | - | - |
| Daccord | - | - | - | - | - | - | - | - |

#### Real sequencing data (Mock communities)

Table 4 shows the error correction benchmarking results for real sequencing data. The three sections of the table show results on the yeast pseudo-diploid genome dataset (mock community) first, the NWC metagenome dataset (real) second, and the Microbial 10-plex metagenome dataset (mock community) as the third section of rows in Table **??**. VeChat achieves approximately 2 ~ 4, 1.4 ~ 7.8 and 3.3 ~ 5.6 times lower error rates on Yeast, NWC and Microbial 10-plex datasets, respectively, while maintaining comparable performance in terms of other aspects.

**Table 4.** Error correction benchmarking results for real sequencing data (mock communities).

| Method | #Seq | Error rate (%) | Mismatch (%) | Indel (%) | Haplotype coverage (%) | N50 (bp) | NGA50 (bp) | # Mis-assemblies |
| --- | --- | --- | --- | --- | --- | --- | --- | --- |
| **Yeast pseudo-diploid genome** |  |  |  |  |  |  |  |  |
| VeChat | 107210 | 0.236 | 0.111 | 0.126 | 99.6 | 5693 | 15537 | 505 |
| Daccord | 149020 | 0.503 | 0.285 | 0.217 | 96.5 | 4762 | 15161 | 1457 |
| Racon | 136199 | 0.758 | 0.282 | 0.476 | 98.3 | 6349 | 16001 | 3836 |
| Canu | 118367 | 0.787 | 0.214 | 0.573 | 99.9 | 5684 | 15603 | 743 |
| CONSENT | 160136 | 0.947 | 0.344 | 0.603 | 99.4 | 5622 | 16001 | 7973 |
| **NWC metagenome** |  |  |  |  |  |  |  |  |
| VeChat | 156426 | 0.101 | 0.031 | 0.070 | 99.3 | 9619 | 27416 | 14961 |
| Daccord | 163313 | 0.140 | 0.079 | 0.061 | 99.5 | 8461 | 26343 | 12440 |
| Racon | 168879 | 0.394 | 0.062 | 0.332 | 99.5 | 9914 | 28428 | 10832 |
| Canu | 37779 | 0.551 | 0.090 | 0.461 | 99.1 | 13811 | 27729 | 4066 |
| CONSENT | 176764 | 0.787 | 0.107 | 0.680 | 99.7 | 9708 | 27638 | 9731 |
| **Microbial 10-plex metagenome** |  |  |  |  |  |  |  |  |
| VeChat | 245804 | 0.089 | 0.066 | 0.023 | 99.3 | 7837 | 17511 | 1533 |
| Racon | 253817 | 0.297 | 0.160 | 0.137 | 97.8 | 8019 | 17760 | 3724 |
| Canu | 193810 | 0.328 | 0.121 | 0.206 | 99.8 | 8477 | 17824 | 1170 |
| Daccord | 254003 | 0.336 | 0.298 | 0.038 | 98.2 | 7704 | 17342 | 2073 |
| CONSENT | 256935 | 0.495 | 0.107 | 0.388 | 100.0 | 8017 | 17729 | 3470 |

### Varying read coverage

In order to evaluate how sequencing coverage influences the error correction approaches, we focuse on the triploid genome, consisting of three *E. coli* strains as described before. We simulated PacBio CLR reads at varying sequencing coverage, namely, 10x, 20x, 30x, 40x, 50x per haplotype.

Supplementary Table S2 shows the benchmarking results of error correction. In summary, VeChat achieves approximately 2 ~ 47 times lower error rates on all datasets, while maintaining better or comparable performance in terms of other aspects such as number of corrected reads, completeness (HC) and length of corrected reads. As the sequencing coverage increases (from 10x to 50x), VeChat achieves better error correction (error rate from 0.311% to 0.017%), while keeping comparable performance in terms of other aspects.

### Varying sequencing error rates

In order to evaluate the effect of sequencing error rate on the different methods, we again focused on the triploid genome consisting of three *E. coli* strains as described above. Accordingly, we simulated PacBio CLR reads at varying sequencing error rates, namely, at 5%, 10% and 15%. The average sequencing coverage per haplotype is 30x.

Supplementary Table S3 shows the corresponding benchmarking results of error correction. Overall, VeChat achieves approximately 10 ~ 93, 9 ~ 26 and 7 ~ 9 times lower error rates on datasets with 5%, 10% and 15% errors, respectively, while maintaining comparable performance in terms of other aspects.

On decreasing sequencing error rate (from 15% down to 5%), VeChat’s error correction undoubtedly improves with the error rate achieved dropping from 0.091% to 0.009%.

### Runtime and memory usage evaluation

The runtime of VeChat is dominated by three steps: while the computation of read overlaps (without base-level alignment) is extremely fast, subsequent edit-distance–based alignment (for segmenting windows) is time-consuming. Second, the POA algorithm that drives the construction of the variation graphs performs sequence to graph alignment, which comes at computational complexity of *O*(*N* (2*N_p_* + 1)|*V* |), where *N* is the length of the sequence to be aligned, *N_p_* is the average number of predecessors in the graph and *V* is the number of vertices in the graph (Lee *et al.*, 2002). Third, VeChat follows an iterative paradigm, such as read overlap computation (with base-level alignment) and error correction (consensus generation) steps during the second iteration; this also requires a considerable amount of running time. We performed all benchmarking analyses on x86 64 GNU/Linux machines using 48 cores. The runtime and peak memory usage evaluations for different methods are reported in Supplementary Tables S4 ~ 7. VeChat takes 23 ~ 81 CPU hours and 8 ~ 92 CPU hours on simulated PacBio CLR and ONT reads from datasets reflecting varying ploidy (2, 3 or 4, as usual), which is 1.1 ~ 6.2 and 0.6 ~ 6.9 times slower than other methods. At the same time, it requires higher peak memory usages (Supplementary Tables S4, S5). VeChat is 2.4 ~ 7.1 times slower than other approaches on the simulated metagenomic dataset, and reaches higher peak memory usages (Supplementary Table S6). Note that the high complexity metagenomic dataset is too large to be processed by Racon and Daccord, whereas our approach is able to handle it. In addition, VeChat is 0.7 32 times slower on real datasets, while requiring higher peak memory usages (Supplementary Table S7).

## Discussion

We have presented VeChat, as an approach that performs haplotype-aware error correction for third-generation sequencing (TGS) reads. To the best of our knowledge, VeChat is the first approach that explicitly addresses to preserve haplotype-specific variation already during the correction process, which improves over prior approaches in particular insofar as bias inducing consensus sequences can be avoided. Results have demonstrated the superiority of VeChat: in all benchmarking scenarios, VeChat suppresses error rates by at least a factor of 2 or 3, if not, as is the case for the majority of scenarios, suppressing error rates by one to two orders of magnitude in comparison with the leading competitors. At the same time, VeChat preserves the haplotype identity of the reads, which means that after correction with VeChat, all reads contribute to the coverage of the haplotype they stemmed from. The most obvious interpretation of these results is that capturing haplotype structure already during error correction is not just beneficial, but perhaps even imperative, when seeking to remove all errors from TGS reads. For appropriately capturing haplotype-specific variants during the error correction step, we construct variation graphs from the noisy TGS reads directly. Note that direct construction of variation graphs from heavily erroneous reads is not standard. In fact, at first glance, it is even counterintuitive, because the seminal idea of variation graphs is to be constructed from true haplotype-specific, sufficiently long patches of sequence. Here, patches of sequence contain up to even 15% of errors.

So, the resulting graph contains a large amount of nodes and edges that reflect such errors. For identifying spurious nodes and edges, one then exploits that sequencing errors are randomly distributed, whereas variants tend to re-occur across different reads. In particular, edges that link spurious nodes (with true nodes or other spurious nodes) tend to be little covered by reads, because reads, unlike for true effects, do not tend to share errors. To systematically identify spurious edges as edges that are covered by too little reads, in a sound, principled way, we adopt two metrics from frequent itemset mining. While Support measures the relative coverage of edges in the graph, Confidence measures the association between the two nodes incident to the edge they share; if basic support or the association is too little, the edge and possibly resulting isolated nodes are removed.

A particular effect of VeChat is to achieve drastic improvements in terms of mismatch rates; improvements on indel rates are also clearly evident in all scenarios, but usually not necessarily drastic. One possible explanation is that substitution events, much more than insertion and deletion events, dominate the evolutionary processes of living organisms, and thus are often characteristic of strains or haplotypes. VeChat appears to be the first approach to correctly preserve these single nucleotide polymorphisms (SNPs), because the distinction of haplotypes is just what variation graphs are made for. At any rate, VeChat appears to prevent masking of true variants as a result of generating consensus sequence.

As for future perspectives, the rapid development of long-read sequencing technologies will lead to decreasing sequencing error rates. However, because the advantages of VeChat become even more evident when sequencing error rates drop, VeChat will also be a superior tool when correcting long reads in which errors appear at a rate of 5% or lower (see Supplementary Table S3).

Of course, future improvements are conceivable: in particular, VeChat requires computational resources that exceed those of other approaches. In particular on large datasets, VeChat experiences longer runtimes and higher peak memory usage. However, there is room for improving on the computational efficiency of VeChat. One crucial point is that VeChat uses off-the-shelf approaches in some places. While these approaches do the job, not all features they provide are needed, which corresponds to an overhead of computations when running the off-the-shelf software. In particular, there is good hope that computations such as edit-distance-based alignments, or the sequence-to-graph alignments, can be replaced by more efficient routines in the future.

## Methods

### Step 1: Read overlap calculation

Step 1 refers to computation of all-vs-all overlaps for the input reads (first cycle: raw reads, second cycle: pre-corrected reads) using the (widely popular) minimizer based, long-read overlap computation tool Minimap2 (Li, 2018). During the first cycle, only a seed-chain procedure is performed, while during the second cycle a base-level alignment is added. Because this is extremely fast, it can easily manage the large amount of read pairs we need to process. Subsequently, bad overlaps are filtered by reasonable, additional criteria, which includes removing overlaps that do not exceed 500 bp in length, self-overlaps, or internal matches. In this, we follow Algorithm 5 in (Li, 2016) and its implementation in (Marijon *et al.*, 2020). Additionally, overlaps that have a high error rate, that is |1 - min(*L_q_, L_t_*)/max(*L_q_, L_t_*)| ≥ *e*, where *L_q_* and *L_t_* are the overlap lengths of the query and target read, respectively, and *e* is the maximum error-rate threshold, are also filtered out.

While during the second cycle, the similar procedures(overlap computation and filtration) are also performed but for pre-corrected reads. Unlike in the first cycle, we compute pre-corrected read overlaps *with base-level alignment* such that the sequence identity of overlaps (overlap identity) can be determined, and then filter overlaps with one more criterion, minimum overlap identity (denoted as *δ*, *δ* = 0.99 for simulated sequencing data and *δ* = 0.98 for real sequencing data). Notably, because most of sequencing errors have been corrected in the first cycle, the identity distribution of read overlaps derived from different haplotypes is clearly dispersed. Therefore, it is straightforward to filter overlaps from different haplotypes by simply setting overlap identity ≥ *δ*.

### Step 2: Read alignment pile generation

Our workflow then selects a target read *r* and, based on the overlaps computed in step 1, collects reads that overlap *r*. The target read *r* serves as a backbone, and for each overlap between read *r* and another read, a fast edit-distance–based alignment (Myers, 1999; Šošić and Šikić, 2017) is then performed, which generates a read alignment pile. The edit-distance–based alignment is only needed to split the read alignment pile into small windows in step 3. Dangling ends of reads that overlap the target read *r*, indicated by horizontal dotted lines in the original read alignment pile in Figure 1, are removed from further consideration in the following. See (Vaser *et al.*, 2017) for more details.

### Step 3: Window segmentation

The read alignment pile is then divided into several small non-overlapping windows of identical length, with the target read serving as a reference: each such window covers 500 bp of the target read. Obviously, segmenting the read alignment pile reflects a straightforward procedure, because of the pairwise alignments of the target read with its overlapping reads served as the basis for read alignment pile construction, see Step 2 (Vaser *et al.*, 2017). The part of the target read *r* corresponding with one particular window is further referred to as a ‘target subread’. This implies in particular that target subreads are 500 bp in length, apart from the rightmost window, where target subreads can be shorter, The reason for segmenting the alignment piles into windows of small length is the great reduction in terms of computational burden in the following: the next step 4, as the technical core of our approach being concerned with variation graphs, greatly profits from this segmentation, both in terms of downsizing the original problem as well as in terms of enabling parallelization.

### Step 4: Error correction for target subreads

Step 4 reflects the methodical novelty of VeChat. Step 4 differs when comparing the first with the second cycle, see Figure 2(a) (a) for the first and Figure 2(a) (b) for the second cycle. Step 4 of the first cycle is considerably more involved, because it reflects the crucial statistical considerations through which to identify sequencing errors. The core idea that underlies these crucial statistical considerations is that true variants are significantly likely to co-occur across different reads, whereas occurrence of errors is random. Correspondingly, the following arguments make sense.

Thanks to dividing reads into sub-reads, the corresponding computations, such as variation graph construction and statistical evaluation of edges relative to read coverage, can be parallelized across sub-reads, which speeds up computations considerably.

#### Variation graph construction

The subsequences in a window we refer to as subreads, are subsequently used to construct a variation graph *G* = (*V, E, P*). This variation graph is a directed acyclic graph (DAG), where vertices *v* ∈ *V* represent nucleotides (A, T, C, G), edges (*v_i_, v_j_*) ∈ *E* indicate that the nucleotides represented by nodes *v_i_* and *v_j_* have appeared as a two-letter subsequence in one of the reads from which the graph was constructed, relative to that particular position with respect to the read alignment pile. So, for example, if *v_i_* and *v_j_* correspond to *A* and *G*, respectively, exactly the reads that relative to the coordinates implied by the target read show *AG* at that particular position induce an edge (*v_i_, v_j_*). Correspondingly, reads can be identified as certain paths *P* = (*v*_1_, *…, v_l_*) in the variation graph. For variation graph construction, we use the partial order alignment (POA) algorithm (Lee *et al.*, 2002) and its faster version, enhanced by SIMD vectorization, as described in (Vaser *et al.*, 2017).

#### Pruning: Principle

In the following, we will use the notation *v ∈ V* to also indicate the letter from the alphabet {*A, C, G, T*} a particular node *v* refers to.

The high error rate affecting TGS reads and the possible bias introduced by the POA algorithm (because, for example, the POA depends on the order relative to which reads are considered), the variation graph constructed in the first cycle contains many spurious vertices and edges.

For pruning the graph from mistaken edges and/or nodes (vertices), we adopt techniques from frequent itemset mining. The basic idea is to identify edges with itemsets, and to prune edges from the graph if the corresponding itemsets do not appear to be sufficiently frequent. After removal of edges, reads are re-aligned against the resulting graph, such that itemset counts have to be re-computed.

This may render more edges to correspond to itemsets that are not sufficiently frequent. The cycle of identifying edges as infrequent itemsets, removing them, and re-aligning reads is repeated until convergence, that is, until no further edges are identified as to be removed. In practice, we determined 3 as an appropriate number of iterations for our experiments. Note that for re-aligning reads against the modified graph, we make use of the POA algorithm, without, however, re-modifying the graph anymore. The model that underlies the mining of frequent itemsets is the “market-basket model”. Baskets correspond to sets of items, and frequent itemsets correspond to subsets of items that appear in sufficiently many baskets, or, vice versa, infrequent itemsets correspond to subsets of items that do not appear in sufficiently many baskets.

Following this model, the basic set of items agrees with the set of nodes *V* in the variation graph. Baskets then correspond to reads, which are modeled as paths *P* = (*v*_1_, *…, v_l_*), and the items they contain correspond to the nodes *v*_1_, *…, v_l_ ∈ V* the reads cover. Further, the itemsets we are interested in correspond to edges *e* = (*v, w*), as pairs of items *v, w*. If two consecutive nodes *v, w*, as particular subsets of items, do not appear in sufficiently many baskets, that is, are not contained in that particular order in sufficiently many paths *P*, the corresponding edge *e* = (*v, w*) is removed from the graph.

For appropriately quantifying “sufficiently many baskets”, we make further use of *Support* and *Confidence* as two standard definitions from frequent itemset mining. While “Support” just corresponds to the number of baskets a particular subset of items is contained in, “Confidence” corresponds to measuring whether appearance of items in basket is correlated (or, in other words, whether sets of items are “associated” with each other). Here, Support just agrees with the number of reads by which a particular edge *e* = (*v, w*) is covered. Confidence corresponds to the amount of reads that cover the edge (*v, w*) in relation to how many reads cover *v*, on the one hand, and in relation to how many reads cover *w*, on the other hand. If neither sufficiently many reads that cover *v* also cover (*v, w*), nor sufficiently many reads that cover *w* also cover (*v, w*), we “loose confidence” in *e* = (*v, w*), because the edge (*v, w*) could reflect sequencing error noise, and remove it.

#### Pruning: Definitions

To make the ideas from above explicit, let *R*(*v*) be all reads that cover node *v*. For *r* ∈ *R*(*v*), let further *p_r,v_* reflect the probability that *v* reflects an error in *r*. When dealing with FASTQ files, the probability *p_r,v_* is derived from the Phred profile of *r*. In case of FASTA files, *p_r,v_* is taken as zero.

We now would like to determine *w*(*v*), as a weight for node *v* that reflects the expected number of reads that cover it. Note that for FASTA files, *w*(*v*) just agrees with the number of reads that cover *v*. For FASTQ files, *w*(*v*) corresponds to summing up 1 − *p_r,v_* across all reads *r ∈ R*(*v*), as the sum of the probabilities that the reads *r ∈ R*(*v*) indeed reflect the letter associated with *v*. In terms of formulas, we obtain

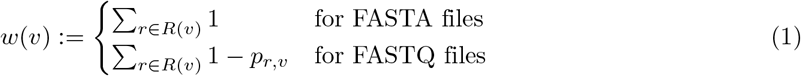

For an edge *e* = (*v_i_, v_j_*), we further determine

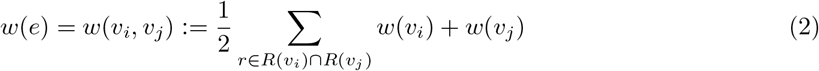

as an approximation for the expected amount of reads that cover *e* = (*v_i_, v_j_*). Note that *w*(*e*) corresponds to exactly the amount of reads that cover *e* for FASTA files. For FASTQ files, the expected amount of reads that cover (*v_i_, v_j_*) virtually corresponds to the sum of products 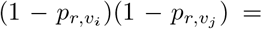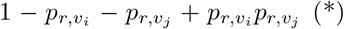 across *r* ∈ *R*(*v_i_*) ∩ *R*(*v_j_*). Here, for speeding up computations, we opt for approximating (*) by 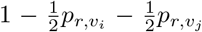. Since this agrees with (*) in terms of orders of magnitude, this introduces only negligible deviations from the true values; experiments of ours (data not shown) confirm that the gain in speed offsets the loss in precision on that account.

Based on these weights, we now define the two metrics *Support* and *Confidence*. See also Figure 3 for an illustration. In formal detail, let *e* = (*v_i_, v_j_*) be an edge. Let then *Support* of *e* be defined as

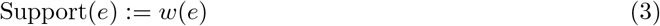

that is, just as the weight of *e*. Further, note that *Confidence* is an asymmetrical measure: the probability to observe *v_j_* in a read that contains *v_i_* may disagree with the probability to observe *v_i_* in a read that contains *v_j_*. We take this into account by defining

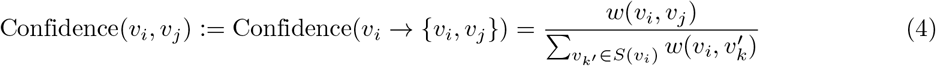

on the one hand, and

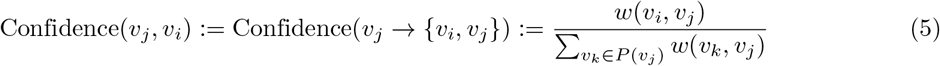

on the other hand, where *S*(*v_i_*) and *P* (*v_j_*) denote the vertices that succeed *v_i_* and precede *v_j_*, respectively (and where *v_i_ → {v_j_, v_j_*} and *v_j_ → {v_i_, v_j_*} agree with standard notation from association rule mining). We eventually declare

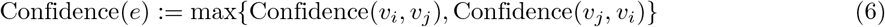

**Figure 3.**
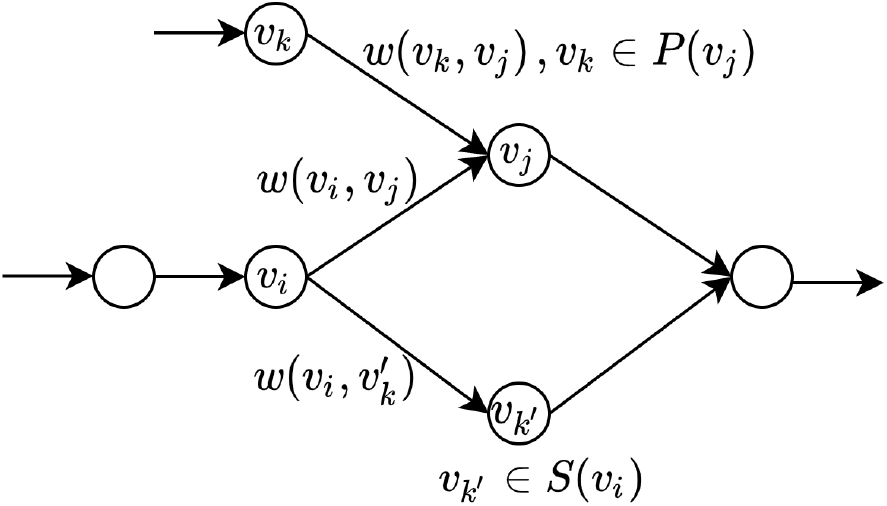
A schematic diagram for explaining the calculation of *Support* and *Confidence* for edge (*v_i_, v_j_*).

Confidence(*e*) ≔ max{Confidence(*v_i_, v_j_*), Confidence(*v_j_, v_i_*)} (6) as the overall Confidence in *e* = (*v_i_, v_j_*).

It remains to determine appropriate thresholds *s* and *c*, such that edges *e* for which either Support(*e*) *< s* or Confidence(*e*):< *c* are pruned from the graph. Note that by its definition, Confidence reflects the probability that a read that covers *v_i_* also covers *v_j_*, or vice versa. We determined *c* = 0.2 as an appropriate threshold in experiments; see the Supplement for the corresponding experiments.

Support, however, does not reflect a probability. Depending on the overall amount of reads in a subwindow, and the length of a subwindow—that is virtually depending on the average read coverage of a subwindow)—the Support needs to be appropriately scaled. Therefore, consider that *C* ≔ Σ_*v ∈ V*_ w(*v*)/*L* is the average coverage of a position in the subwindow. Accordingly, we determine *s* ≔ 0.2 × *C* as a threshold that takes subwindow specific coverage into appropriate account.

Note eventually that both *Support* and *Confidence* are required for effective pruning of the graph, see Figure S2 for the correlation between the two quantities.

#### Optimal alignment path extraction

Upon convergence of the pruning algorithm, the target subread that corresponds to a small window is realigned against the fully pruned variation graph that results from the last iteration of the pruning algorithm. The path in the graph that corresponds to the optimal alignment of the target subread is then taken as the pre-corrected target subread; see the orange elements in Figure 2(a) for an illustration.

### Step 5: Concatenation

In this step, pre-corrected target subreads are concatenated to a whole, pre-corrected target read, which corresponds to the obvious, straightfoward idea of “patching together” pre-corrected target subreads; see “5. Concatenation” in Figure 1 for an illustration.

### Step 6: Merging corrected target reads

Steps 2-5 are repeated until all reads have been corrected. The overall set of pre-corrected that results from these steps are then taken as input for the second cycle.

### Second Cycle: Modifications Step 1 and 4

Note that the pre-corrected reads generated by way of cycle 1 still contain a small, but yet non-negligible amount of random errors. Cycle 2 addresses to correct these remaining errors. To do so, steps 1-6 are repeated with, however, some crucial modifications in steps 1 and 4. See the blue elements in Figure 1 for the workflow that reflects the procedures of cycle 2. To be specific, the modification of step 1 consists in not only computing all-vs-all overlaps based on minimizers, but also *base-level alignments* (Li, 2018) for the pre-corrected reads. This considerably facilitates to filter reads overlaps according to which they stem from identical haplotypes, based on sequence identity related thresholds. We use *δ* = 0.99 for sequencing error rates of 5 10% and *δ* = 0.98 for a sequencing error rate of 15% in our experiments, which we also generally recommend.

The modification of step 4 then relates to *generating a single consensus sequence* from each variation graph instead of performing iterative graph pruning and sequence-to-graph re-alignment. For that, the dynamic programming algorithm (called “heaviest bundle algorithm” because the traversal algorithm selects the path by maximum weight) (Lee, 2003) as indicated in Figure 2(b) is used. Note that generating consensus sequences from graphs is reasonable in the second cycle, because the overlapping reads used to correct the target read stem from the same haplotype. Finally, we obtain fully error-corrected reads as the output of the second cycle; see “All final corrected reads (cycle2)” in Figure 1.

## Acknowledgements

Not applicable.

## Funding

XL and XK were supported by the Chinese Scholarship Council. AS was supported by the Dutch Scientific Organization, through Vidi grant 639.072.309 during the early stages of the project, and from the European Union’s Horizon 2020 research and innovation programme under Marie Skłodowska-Curie grant agreements No 956229 (ALPACA) and No 872539 (PANGAIA).

## Availability of data and materials

Raw sequencing data can be downloaded from Zenodo (DOI: 10.5281/zenodo.5501454). The results can be reproduced from Code Ocean (DOI: 10.24433/CO.2329278.v2). The source code of VeChat is GPL-3.0 licensed, and publicly available at https://github.com/HaploKit/vechat.

## Ethics approval and consent to participate

Not applicable.

## Competing interests

The authors declare that they have no competing interests.

## Consent for publication

Not applicable.

## Authors’ contributions

XL and AS developed the method. XL implemented the software and conducted the data analysis. XK simulated the metagenomic data sets. XL and AS wrote the manuscript. All authors read and approved the final version of the manuscript.

## Notes

### Competing Interest Statement

The authors have declared no competing interest.

