## Supplementary Material for "VeChat: Correcting errors in long reads using variation graphs"

### VeChat: Correcting errors in noisy long reads using variation graphs — Supplementary Material

*To whom correspondence should be addressed.

**Table S1.** Genome descriptions of simulated and real sequencing data sets. See the excel file: *Supplementary Table S1.xlsx*

| Method | #Seq | Error rate (%) | Mismatch (%) | Indel (%) | Haplotype coverage (%) | N50 (bp) | NGA50 (bp) | # Mis-assem-blies |
|---|---|---|---|---|---|---|---|---|
| **Coverage=10x** | | | | | | | | |
| VeChat | 14960 | 0.311 | 0.114 | 0.197 | 99.0 | 12945 | 30952 | 21 |
| Racon | 16308 | 0.629 | 0.433 | 0.196 | 95.8 | 12571 | 30948 | 114 |
| Canu | 16403 | 0.830 | 0.535 | 0.296 | 98.9 | 12584 | 30903 | 35 |
| Daccord | 15896 | 0.851 | 0.749 | 0.102 | 98.5 | 12611 | 31031 | 17 |
| CONSENT | 16802 | 1.075 | 0.708 | 0.367 | 98.4 | 12573 | 30815 | 228 |
| **Coverage=20x** | | | | | | | | |
| VeChat | 31132 | 0.070 | 0.028 | 0.043 | 99.9 | 12765 | 36029 | 15 |
| CONSENT | 33629 | 0.517 | 0.423 | 0.094 | 99.9 | 12559 | 35572 | 121 |
| Racon | 32644 | 0.565 | 0.427 | 0.138 | 97.9 | 12537 | 35503 | 190 |
| Daccord | 32038 | 0.813 | 0.754 | 0.059 | 99.5 | 12550 | 35605 | 12 |
| Canu | 31288 | 0.863 | 0.493 | 0.370 | 99.8 | 12652 | 35581 | 35 |
| **Coverage=30x** | | | | | | | | |
| VeChat | 48085 | 0.031 | 0.015 | 0.016 | 100.0 | 12595 | 38467 | 13 |
| CONSENT | 50462 | 0.276 | 0.205 | 0.071 | 100.0 | 12511 | 38187 | 105 |
| Racon | 48986 | 0.558 | 0.427 | 0.131 | 98.7 | 12484 | 38257 | 288 |
| Canu | 37210 | 0.612 | 0.405 | 0.207 | 99.9 | 13675 | 38360 | 30 |
| Daccord | 48189 | 0.807 | 0.752 | 0.055 | 99.7 | 12485 | 38357 | 16 |
| **Coverage=40x** | | | | | | | | |
| VeChat | 64475 | 0.022 | 0.010 | 0.012 | 100.0 | 12570 | 40264 | 20 |
| CONSENT | 67255 | 0.219 | 0.137 | 0.083 | 100.0 | 12497 | 40077 | 114 |
| Canu | 50236 | 0.533 | 0.334 | 0.199 | 100.0 | 13678 | 39999 | 30 |
| Racon | 65210 | 0.561 | 0.425 | 0.136 | 99.1 | 12482 | 39925 | 386 |
| Daccord | 64294 | 0.803 | 0.749 | 0.054 | 99.7 | 12466 | 39970 | 17 |
| **Coverage=50x** | | | | | | | | |
| VeChat | 80984 | 0.017 | 0.008 | 0.009 | 100.0 | 12537 | 41609 | 34 |
| CONSENT | 84185 | 0.211 | 0.112 | 0.099 | 100.0 | 12477 | 41284 | 124 |
| Canu | 48663 | 0.459 | 0.295 | 0.164 | 100.0 | 14427 | 41217 | 28 |
| Daccord | 80529 | 0.804 | 0.751 | 0.053 | 99.8 | 12428 | 41311 | 25 |

**Table S2.** Error correction benchmarking results for simulated PacBio CLR reads of various sequencing coverages. The average sequencing coverage per haplotype is set as 10x, 20x, 30x, 40x, and 50x, respectively. The ploidy is three and sequencing error rate is 10%.

| Method | #Seq | Error rate (%) | Mismatch (%) | Indel (%) | Haplotype coverage (%) | N50 (bp) | NGA50 (bp) | # Mis-assem-blies |
|---|---|---|---|---|---|---|---|---|
| **Error=5%** | | | | | | | | |
| VeChat | 48624 | 0.009 | 0.007 | 0.003 | 100.0 | 12764 | 38945 | 19 |
| CONSENT | 50144 | 0.091 | 0.073 | 0.018 | 100.0 | 12724 | 38779 | 80 |
| Racon | 48691 | 0.507 | 0.440 | 0.067 | 98.2 | 12724 | 38257 | 497 |
| Canu | 35914 | 0.539 | 0.460 | 0.079 | 99.8 | 13996 | 38468 | 116 |
| Daccord | 47645 | 0.838 | 0.784 | 0.054 | 99.7 | 12705 | 38389 | 82 |
| **Error=10%** | | | | | | | | |
| VeChat | 48085 | 0.031 | 0.015 | 0.016 | 100.0 | 12595 | 38467 | 13 |
| CONSENT | 50462 | 0.276 | 0.205 | 0.071 | 100.0 | 12511 | 38187 | 105 |
| Racon | 48986 | 0.558 | 0.427 | 0.131 | 98.7 | 12484 | 38257 | 288 |
| Canu | 37210 | 0.612 | 0.405 | 0.207 | 99.9 | 13675 | 38360 | 30 |
| Daccord | 48189 | 0.807 | 0.752 | 0.055 | 99.7 | 12485 | 38357 | 16 |
| **Error=15%** | | | | | | | | |
| VeChat | 46829 | 0.091 | 0.049 | 0.042 | 99.9 | 12650 | 36292 | 67 |
| Racon | 48359 | 0.618 | 0.390 | 0.228 | 99.7 | 12539 | 36241 | 163 |
| CONSENT | 50032 | 0.680 | 0.429 | 0.252 | 99.9 | 12532 | 36054 | 218 |
| Daccord | 47914 | 0.780 | 0.716 | 0.064 | 99.8 | 12527 | 36338 | 5 |
| Canu | 39040 | 0.843 | 0.364 | 0.479 | 99.9 | 13333 | 36045 | 28 |

**Table S3.** Error correction benchmarking results for simulated PacBio CLR reads with different sequencing error rates of polyploid genome (ploidy=3). The average sequencing coverage per haplotype is 30x and sequencing error rate = 5%, 10%, 15%.

| **Method** | **CPU time (h)** | **Peak memory usage (GB)** |
|---|---|---|
| **Ploidy=2** | | |
| CONSENT | 4.8 | 5.3 |
| Daccord | 12.7 | 12.8 |
| Canu | 14.7 | 5.0 |
| Racon | 17.9 | 4.2 |
| VeChat | 23.3 | 10.8 |
| **Ploidy=3** | | |
| CONSENT | 8.7 | 7.8 |
| Canu | 14.6 | 5.1 |
| Daccord | 26.0 | 14.0 |
| Racon | 39.0 | 7.3 |
| VeChat | 47.1 | 24.1 |
| **Ploidy=4** | | |
| CONSENT | 13.1 | 8.8 |
| Canu | 22.6 | 5.1 |
| Daccord | 51.7 | 17.1 |
| Racon | 77.2 | 10.7 |
| VeChat | 81.4 | 29.8 |

**Table S4.** Runtime and memory usage for simulated PacBio CLR reads of various polyploid genomes (ploidy=2,3,4). The average sequencing coverage per haplotype is 30x and sequencing error rate is 10%.

| Method | CPU time (h) | Peak memory usage (GB) |
|---|---|---|
| **Ploidy=2** | | |
| CONSENT | 4.9 | 5.1 |
| Daccord | 8.4 | 12.9 |
| Canu | 11.6 | 5.0 |
| Racon | 14.2 | 5.3 |
| VeChat | 8.4 | 28.3 |
| **Ploidy=3** | | |
| CONSENT | 8.5 | 7.5 |
| Canu | 11.9 | 5.2 |
| Daccord | 17.4 | 14.0 |
| Racon | 31.2 | 7.7 |
| VeChat | 46.4 | 50.5 |
| **Ploidy=4** | | |
| CONSENT | 13.4 | 8.8 |
| Canu | 28.0 | 5.0 |
| Daccord | 30.9 | 17.1 |
| Racon | 35.6 | 11.7 |
| VeChat | 91.8 | 54.4 |

**Table S5.** Runtime and memory usage for simulated Oxford Nanopore reads of various polyploid genomes (ploidy=2,3,4). The average sequencing coverage per haplotype is 30x and sequencing error rate is 10%.

| Method | CPU time (h) | Peak memory usage (GB) |
|---|---|---|
| **Low complexity (20 genomes)** | | |
| CONSENT | 56.5 | 15.8 |
| Canu | 100.8 | 10.8 |
| Racon | 159.3 | 44.2 |
| Daccord | 162.9 | 8.9 |
| VeChat | 393.5 | 51.0 |
| **High complexity (100 genomes)** | | |
| CONSENT | 375.0 | 17.6 |
| Canu | 880.0 | 10.8 |
| VeChat | 2668.7 | 78.0 |
| Racon | - | - |
| Daccord | - | - |

**Table S6.** Runtime and memory usage for simulated PacBio CLR reads of metagenomic datasets with different complexity. The average sequencing coverage of strains is 30x and sequencing error rate is 10%.

| Method | CPU time (h) | Peak memory usage (GB) |
|---|---|---|
| **Yeast pseudo-diploid genome** | | |
| CONSENT | 5.2 | 14.0 |
| Canu | 29.1 | 5.2 |
| Racon | 33.2 | 8.3 |
| Daccord | 77.7 | 16.5 |
| VeChat | 51.5 | 42.4 |
| **NWC metagenome** | | |
| CONSENT | 24.4 | 19.7 |
| Canu | 26.2 | 5.0 |
| Daccord | 208.6 | 15.3 |
| Racon | 423.1 | 25.5 |
| VeChat | 780.0 | 61.6 |
| **Microbial 10-plex metagenome** | | |
| CONSENT | 37.4 | 13.2 |
| Canu | 26.2 | 5.0 |
| Daccord | 80.8 | 11.8 |
| Racon | 152.7 | 29.4 |
| VeChat | 285.9 | 40.8 |

**Table S7.** Runtime and memory usage for real sequencing data (mock communities).

**(a)**

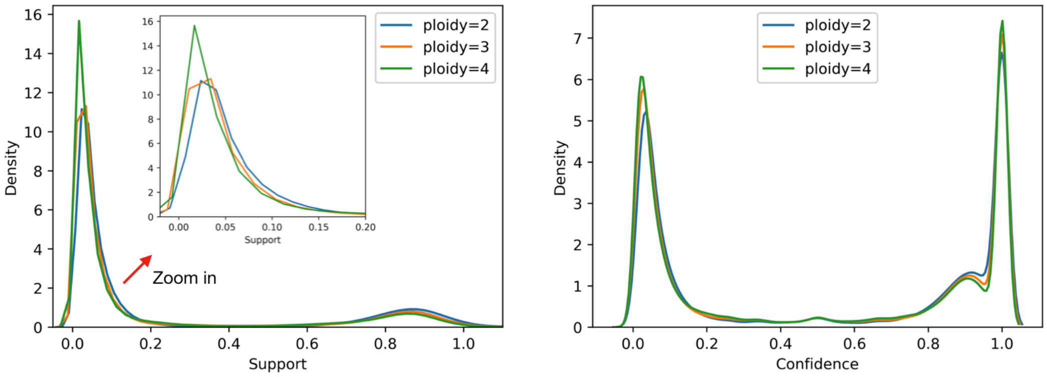

Density
Support
Zoom in
ploidy=2
ploidy=3
ploidy=4
Confidence


**(b)**

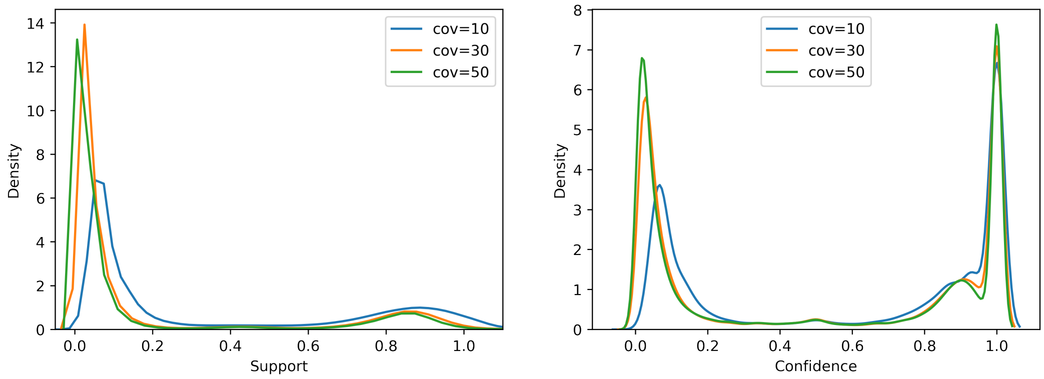

Density
Support
cov=10
cov=30
cov=50
Confidence


**(c)**

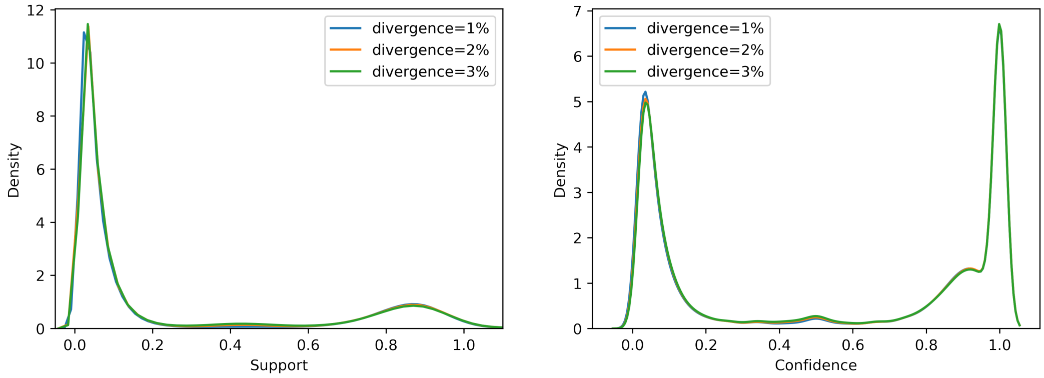

Density
Support
divergence=1%
divergence=2%
divergence=3%
Confidence

**(d)**

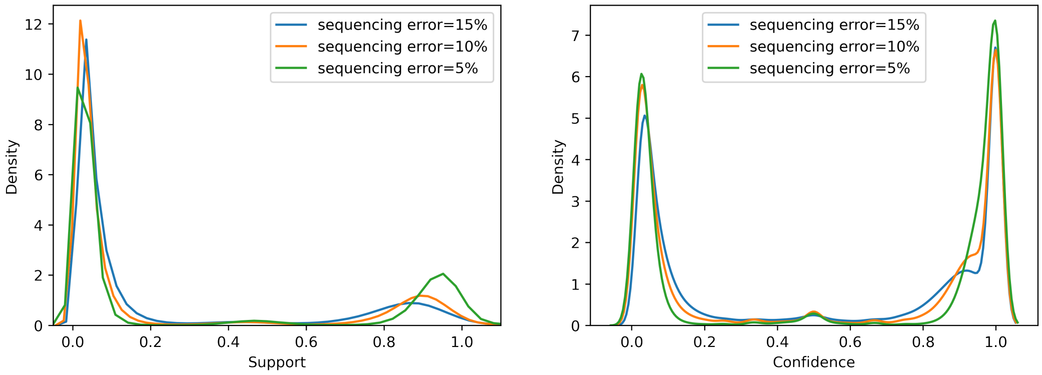


**(e)**

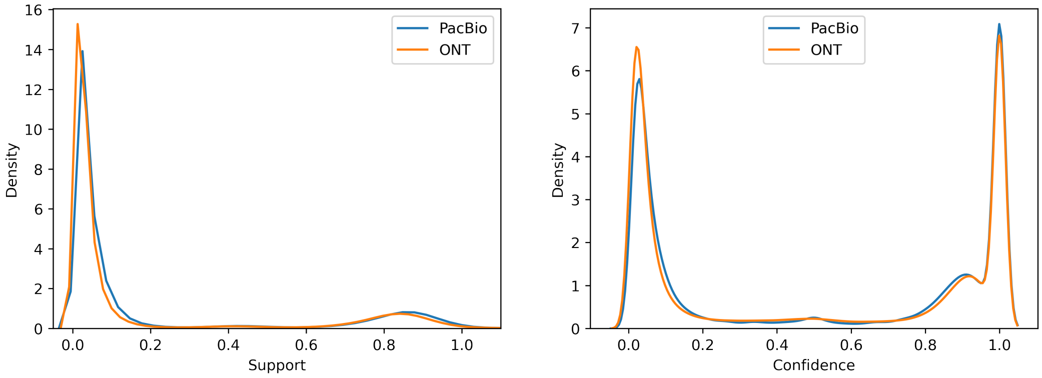


**Figure S1.** Distributions for *Support* and *Confidence* in the raw variation graph in various settings. **(a), (b), (c), (d), (e)** show the different settings for ploidy, sequencing coverage, genome divergence, sequencing error rate and sequencing platform, respectively. The original genome sequence (length=50Kbp) used in this experiment is generated by randomly choosing $\{A, T, C, G\}$, and the mutated haplotypes are generated by introducing random mutations artificially.

**(a)**

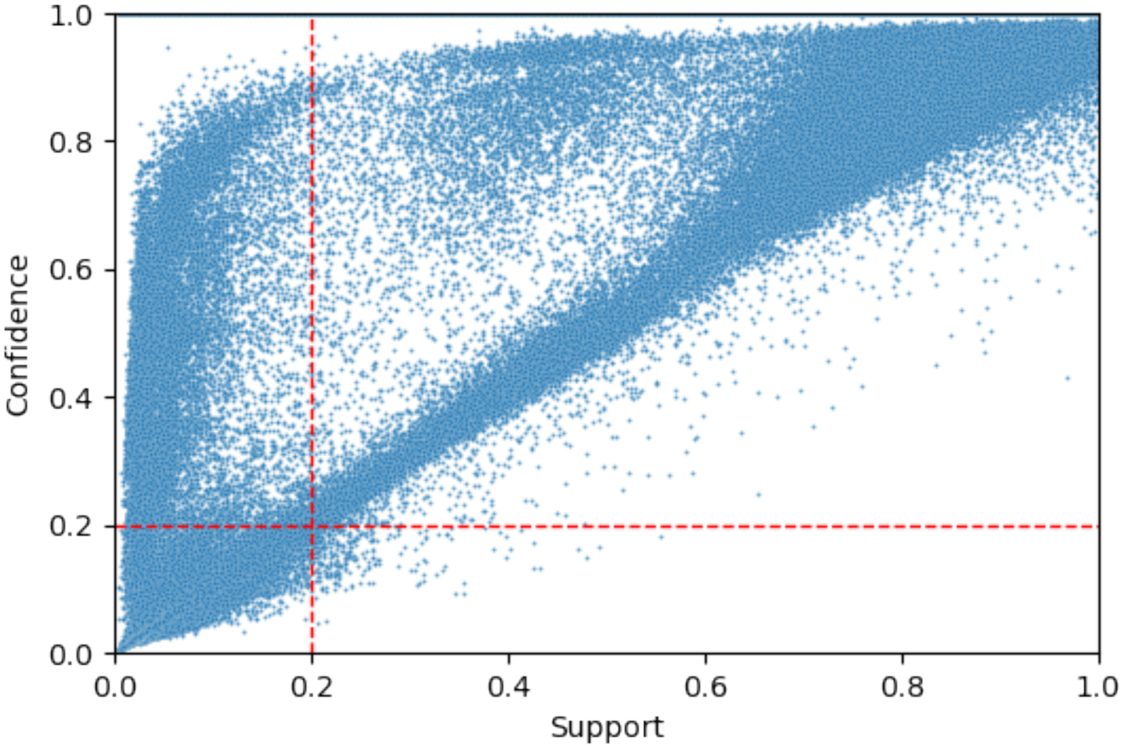


**Figure S2.** *Support* and *Confidence* in the raw variation graph. The dataset is from a simulated diploid genome with 2% genome divergence and 50Kbp genome length. The genome sequence is generated randomly and sequencing coverage 20x per haplotype. The reads are simulated PacBio CLR reads with error rate 15%. Each point in the figure represents an edge. The red dashed lines indicate the thresholds of *Support* and *Confidence* that used in our experiments. Only the points (edges) on the top right region are retained after graph pruning process.

#### Commands and versions of tools used for comparison

- PBSIM2
  `pbsim2 --accuracy-mean 0.9 --hmm_model P6C4.model $ref` #PacBio
  `pbsim2 --accuracy-mean 0.9 --hmm_model R103.model $ref --difference-ratio 23:31:46` #ONT

- Canu v2.1.1
  `canu genomeSize=$genomesize -pacbio $raw_read`
  `canu genomeSize=$genomesize -nanopore $raw_read`
  `canu genomeSize=$genomesize -pacbio-hifi $corrected_read` #run HiFi mode on corrected reads

- Flye v2.8.2-b1689
  `flye --pacbio-raw  $read` #(PacBio)
  `flye --nano-raw  $read` #(ONT)
  `flye --meta --pacbio-raw  $read` #(metagenome assembly)

- Racon v1.4.13
  `racon -f  $raw_read  $overlap  $raw_read >$corrected_read`

- Daccord v0.0.18
  `fasta2DAM reads.dam reads`
  `DBsplit -s256 -x1000 reads.dam`
  `HPC.daligner reads.dam -T$threads| bash`
  `daccord reads.las reads.dam >$corrected_read`

- CONSENT v2.2.2
  `CONSENT-correct --in $raw_read --out $corrected_read  --type PB/ONT`

- QUAST v5.1.0rc1
  `quast.py -r $ref  --min-contig 500 -o out --ambiguity-usage one --fast $fa`

- VeChat v1.1.0
  #(simulated data)
  `vechat -o $corrected_read --platform pb/ont  $raw_read`

  #(simulated data, metagenomic dataset of high complexity)
  `vechat -o $corrected_read --platform pb/ont  --split --split-size 100000 $raw_read`

  # (real data: Yeast pseudo-diploid/Microbial 10-plex metagenome)
  `vechat -o $corrected_read --platform pb --min-identity-cns 0.98 --scrub $raw_read`

  # (real data: NWC)
  `vechat -o $corrected_read --platform pb --min-identity-cns 0.98`
  `--split --split-size 70000 $raw_read`

- VeChat + HiCanu
  `canu genomeSize=$genomesize  -pacbio-hifi $corrected_read`

- VeChat + Canu
  `canu genomeSize=$genomesize  -corrected -pacbio $corrected_read`

- VeChat + Flye
  `flye  --pacbio-corr  $corrected_read`
  `flye --meta  --pacbio-corr  $corrected_read` #(metagenome)

- minimap2 v2.17

- fpa v0.5

- yacrd v0.6.2
